# Collapsing retroviruses for efficient delivery of viro-toxic cargoes

**DOI:** 10.64898/2026.06.28.735095

**Authors:** Christopher D. Mullally, Bojana Stefanovska, Yanjun Chen, Harshita B. Gupta, Michael A. Carpenter, Reuben S. Harris

## Abstract

Retroviruses are excellent tools for delivering and expressing transgenic cargoes with broad utility in research and therapy. However, many cargoes including virus restriction factors can dramatically limit virus production and/or infectivity. An extreme viro-toxic cargo is the DNA cytosine deaminase APOBEC3B, which potently restricts retrovirus infectivity by a direct cDNA deamination-dependent mechanism. To overcome viro-toxicity, an APOBEC3B minigene cargo is disrupted by a translation stop cassette flanked by a direct repeat of its own sequence. The integrity of the minigene is reconstituted naturally by retroviral recombination during transduction. Efficiency can be improved from 90% to nearly 100% by coupling the minigene to the translation of a downstream selectable marker. Collapsing retrovirus (CRV) technology enables the functional delivery of APOBEC3B to a target cell population and may have broad utility for delivering viro-toxic cargoes.

**Motivation:** Many different proteins are detrimental to retrovirus production and/or to virus-producing cells. This problem can be circumvented by interrupting the genetic cargo with a direct repeat-flanked translation stop cassette and allowing transgene integrity to be restored naturally by recombination during reverse transcription. As exemplified by the retrovirus restriction factor APOBEC3B, this process is very efficient.

**Highlights:**

- Collapsing retroviruses (CRVs) prevent retrovirus restriction by viro-toxic proteins
- Truncated tandem repeats recombine during transduction and restore virus integrity
- Translation-coupled selection enables near 100% transduction efficiency
- CRVs have broad utility for delivering viro-toxic cargoes such as APOBEC3B

**Graphical abstract:** 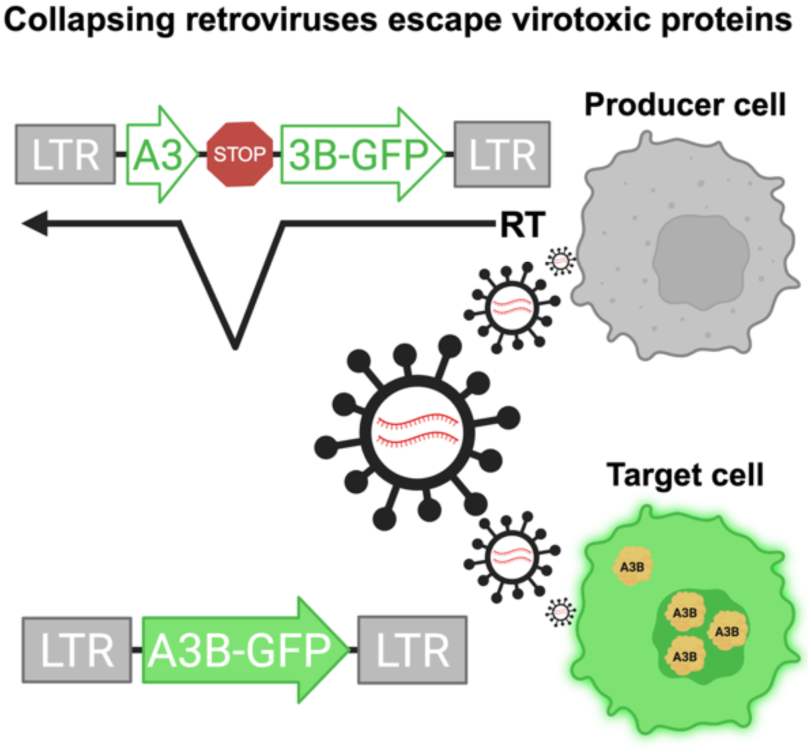

## Introduction

Retroviral transduction is an established method to deliver a genetic cargo to a target cell population. Cargoes range from protein coding cDNA sequences for complementation and over-expression studies to non-coding RNAs for genetic knockdowns or knockout experiments.

Retroviruses are additionally useful because they can deliver relatively large cargoes up to approximately 19 kbp (Shin et al., 2000). A specific example of the utility of retrovirus transduction is delivery of a cytosine base editing (CBE) complex, which is comprised of an APOBEC family DNA deaminase domain fused to a Cas9-nickase-Ugi complex (along with a separately encoded guide RNA), to a target cell population for efficient mutation of a single target cytosine (converting C/G to T/A) (Osgood et al., 2025; Rallapalli and Komor, 2023). This application has enormous promise for multiple applications including gene therapy, where the detrimental effects of inherited or somatic mutations can be reversed through single, precisely targeted C/G to T/A mutations (Coelho et al., 2024; Rees and Liu, 2018).

One limitation to retrovirus transduction technologies is toxicity of the cargo to the virus itself (*i.e*., viro-toxicity). Some factors, if expressed in virus producing cells, can dominantly interfere with virus assembly, budding, titer, and/or infectivity. These viro-toxic factors include a class of proteins that have evolved to be natural antagonists of virus replication. Examples of retrovirus restriction factors include TETHERIN, SAMHD1, and the APOBEC family of DNA cytosine deaminases (Colomer-Lluch et al., 2018; Harris and Dudley, 2015; Wolf and Goff, 2008). APOBEC3G (A3G), for instance, is a potent retrovirus restriction factor that packages into assembling viral particles and through deamination-dependent and independent mechanisms destroys the viral single-stranded cDNA during reverse transcription (Harris et al., 2003; Mangeat et al., 2003; Newman et al., 2005; Schumacher et al., 2008; Zhang et al., 2003). A hallmark of A3G-mediated restriction is >1000-fold increase in viral (genomic strand) G-to-A hypermutation, which is the direct product viral cDNA C-to-U deamination (Harris *et al*., 2003). The HIV-1 accessory protein Vif functions naturally to protect the virus by degrading A3G in virus-producing cells but this mechanism is not completely efficient, especially if A3G is overexpressed is *cis* or *trans* to the viral genome itself in the virus-producing cells.

In addition to being virus restriction factors, at least two APOBEC3 family members, APOBEC3A (A3A) and APOBEC3B (A3B), are also potent cancer mutagens (Carpenter et al., 2023; Mertz et al., 2022; Petljak et al., 2022). Aberrant or dysregulated expression of A3A and A3B results in genomic DNA deamination and signature single base substitution mutations in nearly 70% of human cancer types and is the dominant mutagen in several including tumors of the bladder, breast, cervix, lung, head/neck tissues. Studies here were motivated by the fact that A3B is equally potent as A3G in restricting retrovirus infectivity (Doehle et al., 2005; Hultquist et al., 2011), which complicates the utilization of current retrovirus transduction technologies to deliver catalytically active A3B constructs to difficult-to-transfect normal and cancer cell populations. Inspired by an earlier study (Delviks-Frankenberry et al., 2019), a panel of collapsing retrovirus (CRV) constructs was developed to be able to use retrovirus transduction technology to delivery A3B to target cell populations efficiently and with remarkably high-fidelity. Together with other technologies including 2A translation read-through sequences, drug selection, and anti-retroviral compounds, the efficiency and fidelity of functional A3B delivery to a target population can approach 100%.

## Results

### Collapsing retrovirus (CRV) technology prevents restriction by A3B

To overcome the retrovirus restriction barrier imposed by A3B, a murine leukemia virus (MuLV)-based construct was assembled in which an *A3B* minigene was split into two non-functional parts, each with 1030 bp of self-homology (see schematic in **Figure 1A** and a more detailed illustration in **Figure S1**). This construct includes a 133 bp chimeric intron to prevent A3B toxicity in *E. coli* during plasmid preparation, and a translation stop cassette containing three in-frame stop codons to prevent leaky expression in virus-producing cells. These modifications are implemented to prevent expression of a catalytically active form of A3B in the virus-producing cells. However, following transduction of a target cell population, reverse transcription-mediated template switching causes the nascent viral cDNA to skip from one direct repeat to the other, which leads to an in-frame recombination event between the two homologous regions. The net effect is the seamless collapse of the direct repeats into a single, non-repetitive sequence, which restores functionality to the *A3B* minigene upon the completion of reverse transcription and integration into target cell genomic DNA (see schematic in **Figure 1A** and a step-by-step illustration in **Figure S1**). Henceforth, this approach is referred to as collapsing retrovirus technology (CRV1.0, 2.0, *etc*.). The CRV1.0 constructs also have a downstream internal ribosome entry sequence (IRES) and drug resistance cassette (puromycin resistance) to facilitate selection of transduced cell populations.

**Figure 1.**
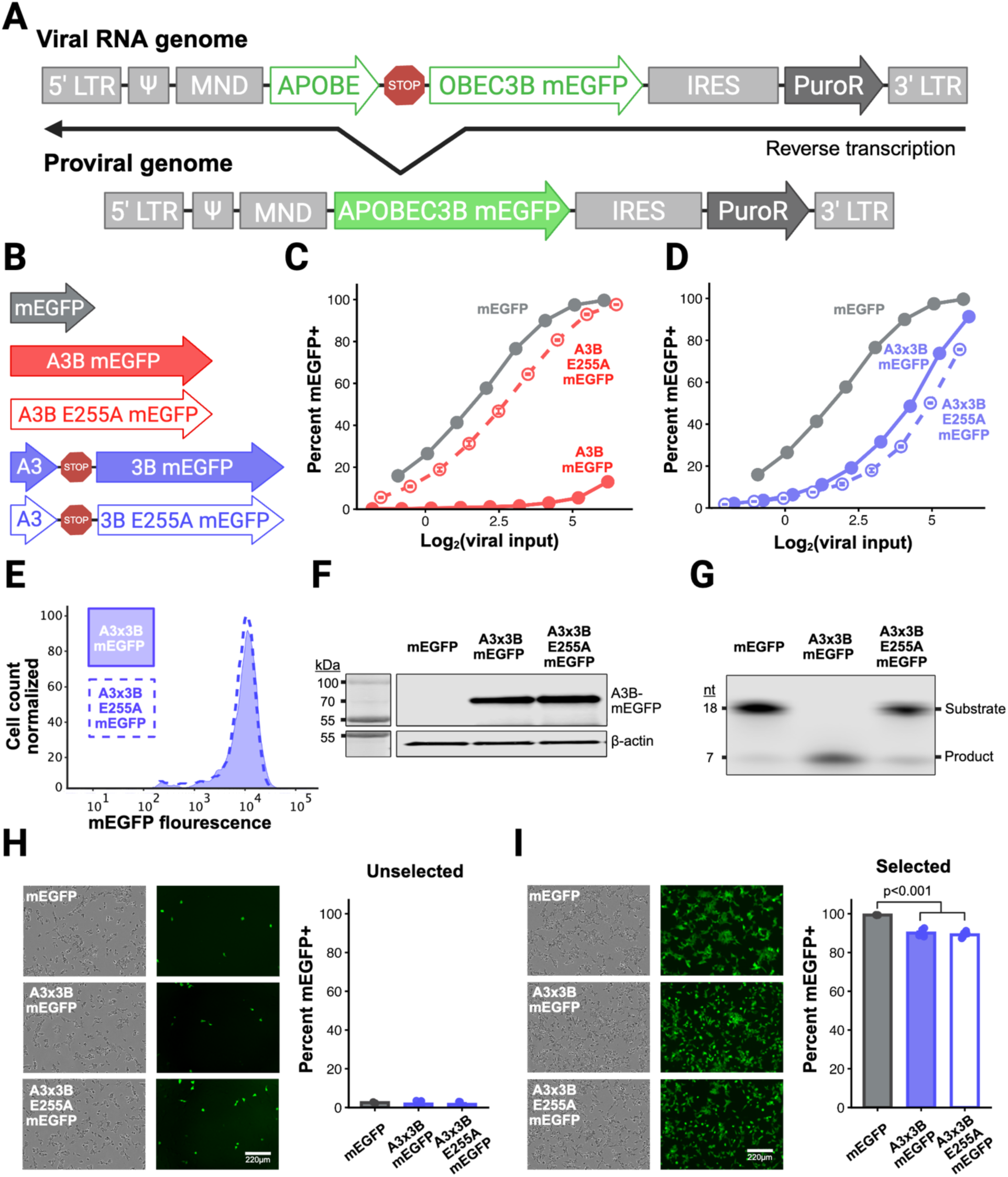
CRV1.0: Collapsing retroviruses enable efficient A3B minigene delivery. **A**, Illustration of mLV-based collapsing retroviruses recombining during reverse transcription in CRV1.0 vectors. MND is a chimeric viral constative promotor. IRES is internal ribosome entry site. PuroR is an open-reading frame encoding puromycin n-acetyl transferase. **B**, Schematic of the panel of CRV1.0 constructs being tested. **C-D**, Infectivity of non-collapsing retroviruses and otherwise isogenic CRV1.0 constructs 72 hours after transduction with the indicated range of virus stocks [mEGFP (gray), A3B-mEGFP (solid red), A3B-E255A-mEGFP (dashed red), A3×3B-mEGFP (solid blue), A3×3B-E255A-mEGFP (dashed blue)]. Viral input measured in pg of viral genomes by RT-qPCR. Data represent the mean of three biological replicates ± SEM with the error bars smaller than datapoints in some cases. **E**, Histogram of mEGFP mean fluorescent intensity measured by live-cell flow cytometry 293T cells transduced with CRV1.0 A3×3B-mEGFP (solid blue) and A3×3B-E255A-mEGFP (dashed blue). **F**, Immunoblot of 293T cellular lysates with CRV1.0 minigenes expressing mEGFP, A3B-mEGFP, or A3B-E255A-mEGFP. **G**, *In vitro* deaminase cleavage assay of 18mer hairpin oligo incubated with 293T cellular lysates from cells transduced with CRV1.0 expressing mEGFP, A3B-mEGFP, or A3B-E255A-mEGFP. **H**-**I**, Representative images and quantification of percent mEGFP+ of 293T transduced with CRV1.0 mEGFP, A3B-mEGFP, or A3B-E255A-mEGFP pre- and post-selection. Data represent the mean of three biological replicates ± SEM with individual datapoints jittered.

To test whether CRV1.0 constructs can overcome A3B-mediated retrovirus restriction, vector matched controls were generated with monomeric enhanced mEGFP (mE, monomeric enhanced; GFP, green fluorescent protein), A3Bi-mEGFP, catalytically inactive A3Bi-E255A-mEGFP for comparison with the collapsing retrovirus counterparts A3×3Bi-mEGFP and A3×3Bi-E255A-mEGFP (i, intron; E255A, catalytic glutamate to alanine mutant; **Figure 1B**). As anticipated from prior studies with A3B expressed in *trans* to the retrovirus (Doehle *et al*., 2005; Hultquist *et al*., 2011), expression of A3Bi-mEGFP in *cis* from the proviral plasmid DNA itself potently restricts the infection of the emergent viral particles (**Figure 1C**). Moreover, MuLV restriction is largely dependent on A3B catalytic activity, as an otherwise isogenic catalytic mutant construct (E255A) is nearly as infectious as a smaller-sized mEGFP control construct (**Figure 1C**). In contrast, the infectivity of the CRV1.0 A3×3Bi-mEGFP construct, as quantified by mEGFP fluorescence of the transduced 293T target cell population, is similar to that of the CRV1.0 A3×3Bi-E255A-mEGFP catalytic mutant construct (**Figure 1D**). Importantly, the CRV1.0 constructs enabled both active A3B and catalytic mutant A3B to be expressed at similar levels in target cells by flow cytometry and immunoblotting, and wildtype A3B but not A3B-E255A is able to catalyze C-to-U deamination of a single-stranded DNA substrate in 293T whole cell extracts above background levels (**Figure 1E, F, G**). Last, but not least, selection for drug-resistance results in approximately 90% of transduced target cells expressing A3B or A3B-E255A (90.1% and 89.1%, respectively), which is a relatively high efficiency but still significantly lower than the 99.3% observed with the mEGFP control (**Figure 1H, I**).

### Mechanistically linking recombination and drug selection improves CRV efficiency

Despite a high recombination rate, the IRES-based CRV1.0 exhibited a significant drawback. Recombination quantified by mEGFP fluorescence appeared to plateau around 90%, which leaves a subpopulation of cells not expressing A3B. Moreover, as overexpressed A3B is known to be genotoxic, transduced target cells that are not expressing active A3B may be able to expand preferentially and overtake the desired A3B-expressing population. The non-fluorescent cells are unlikely due to a poor drug-resistance selection, as parallel mock-transduced 293T cells were killed completely by puromycin.

To address this issue, the constructs were modified to replace the IRES with tandem 2A ribosome-skipping peptide tags (P2A-T2A; **Figure 2A, B**). This CRV2.0 strategy ensured that expression of the puromycin N-acetyltransferase resistance cassette is now dependent upon an in-frame (and therefore more likely to be error-free) recombination event joining the homologous regions of the *A3B* minigene. Two 2A sequences were used to help ensure high rates of ribosome skipping and minimal chimeric A3B-PuroR translation products.

**Figure 2.**
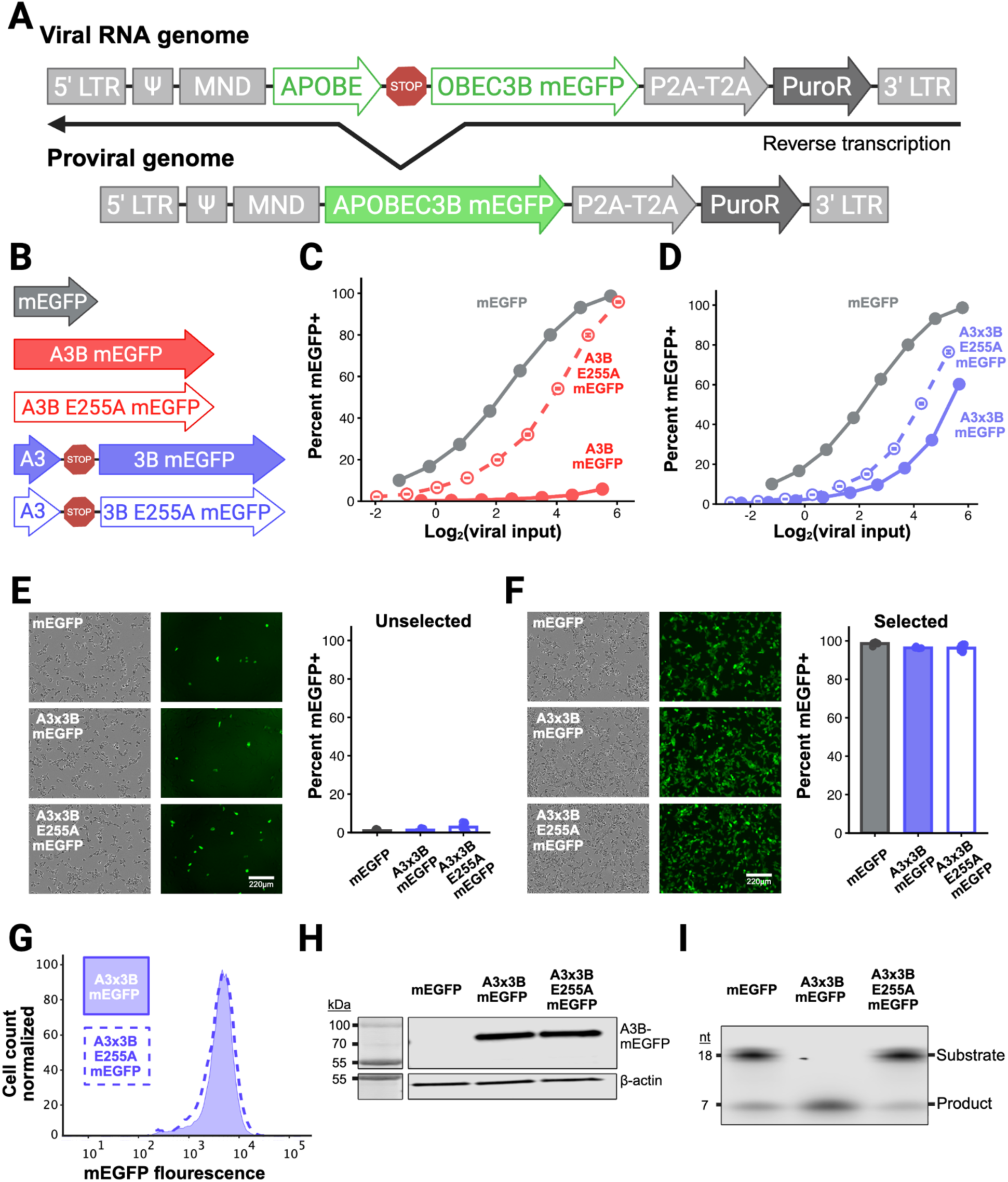
CRV2.0: Recombination-dependent selection creates pools uniformly expressing A3B. **A**, Illustration of mLV-based collapsing retroviruses recombining during reverse transcription in CRV2.0 vectors. MND is a chimeric viral constative promotor. P2A-T2A are tandem viral “self-cleaving” peptide sequences. PuroR: open-reading frame encoding puromycin n-acetyl transferase. **B**, Schematic of the panel of CRV2.0 constructs being tested. **C-D**, Infectivity of non-collapsing retroviruses and otherwise isogenic CRV2.0 constructs 72 hours after transduction with the indicated range of virus stocks [mEGFP (gray), A3B-mEGFP (solid red), A3B-E255A-mEGFP (dashed red), A3×3B-mEGFP (solid blue), A3×3B-E255A-mEGFP (dashed blue)]. Viral input measured in pg of viral genomes by RT-qPCR. Data represent the mean of three biological replicates ± SEM with the error bars smaller than datapoints in some cases. **E**-**F**, Representative images and quantification of percent mEGFP+ of 293T cells transduced with CRV2.0 mEGFP, A3B-mEGFP, or A3B-E255A-mEGFP pre- and post-selection. Data represent mean of three biological replicates ± SEM with individual datapoints jittered. **G**, Histogram of mEGFP mean fluorescent intensity measured by live-cell flow cytometry 293T cells transduced with CRV2.0 A3×3B (solid blue) or A3×3B E255A (dashed blue). **H**, Immunoblot of 293T cellular lysates with CRV2.0 minigenes expressing mEGFP, A3B-mEGFP, orA3B-E255A-mEGFP. **I**, *In vitro* deaminase cleavage assay of 18mer hairpin oligo incubated with 293T cellular lysates from cells transduced with CRV2.0 expressing mEGFP, A3B-mEGFP, or A3B-E255A-mEGFP.

With these CRV2.0 constructs, a new serial dilution viral titration experiment was done, which reinforced the findings above and showed that A3×3B collapsing retroviruses significantly rescues viral infectivity (**Figure 2C, D**). Importantly, the P2A/T2A-mediated selection strategy improves recovery of the A3×3Bi-mEGFP and A3×3Bi-E255A-mEGFP transduced populations to nearly 100% (98.6% mEGFP, 96.4% A3×3Bi-mEGFP, and 96.3% A3×3Bi-255A-mEGFP, respectively (**Figure 2E, F**). As above, live cell flow cytometry and immunoblots show that A3B-mEGFP and A3B-255A-mEGFP are expressed at similar levels in the selected 293T target cell population, but only the former protein catalyzes DNA deamination in whole cell extracts above background levels (**Figure 2G, H, I**). These results combine to indicate that the collapsing retrovirus strategy for transducing viro-toxic proteins such as A3B can be engineered to be highly efficient.

### Retrovirus mutagenesis by A3B is significantly attenuated by CRV2.0 strategy

A hallmark of A3B-dependent retrovirus restriction is cDNA cytosine to uracil deamination, which results in genomic strand G-to-A hypermutation. To assess the genetic integrity of integrated CRV2.0 constructs, proviruses were PCR amplified from the genomes of HeLa cells infected with standard and CRV constructs and subjected to Oxford Nanopore long-read sequencing (**Figure 3A** with supporting data in **Figure S2**). Transduced cells were unselected allowing for an unbiased assessment of recombination rates. Approximately 90% of the full length reads map to seamlessly collapsed retroviral constructs. The remaining ∼10% (A3×3B-mEGFP 11.2%, A3×3B-E255A-mEGFP 8.7%) retained the full tandem repeat cassette, which reflects the experimentally estimated recombination rate above in Figure 1 (**Figure 3B**). Moreover, the frequency of insertion-deletion mutations (indels) was low and similar between all constructs (*i.e.,* both non-collapsing and collapsing CRV integrants) suggesting that homology-mediated recombination during reverse transcription is high fidelity and does not lead to increased indel frequency (**Figure 3C**). Not surprisingly, integrants obtained from the active A3B-mEGFP constructs (non-CRV) in Figure 1 showed very high frequencies of G-to-A mutations, with 33.2 ± 1.7 APOBEC signature G-to-A mutations per full-length viral genome (**Figure 3D**). Importantly, the CRV2.0 A3×3B-mEGFP condition enabled a substantially attenuated mutation burden of 8.9 ± 3.1 APOBEC signature G-to-A mutations viral genome (**Figure 3D**).

**Figure 3.**
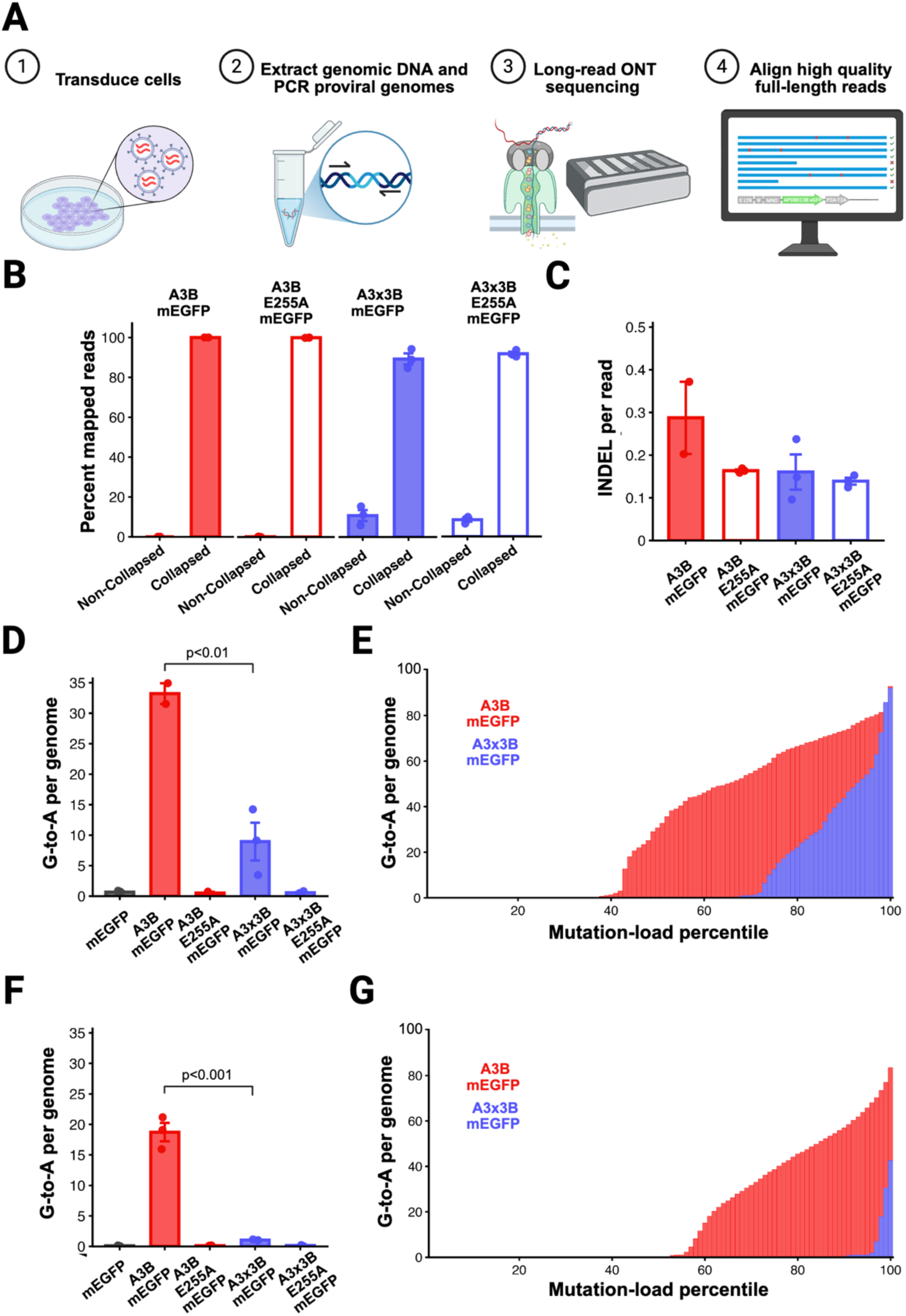
Sequencing indicates CRV proviral insertions are high-fidelity. **A**, Representative workflow. HeLa and 293T were transduced with CRV2.0 constructs followed by genomic DNA isolation, PCR of integrated proviral genomes, ONT long-read sequencing, and alignment of full-length reads. **B**, Percentage of full-length reads mapping exclusively to collapsed or non-collapsed references. Data represent the mean of three biological replicates ± SEM with individual datapoints jittered. **C**, Number of insertion/deletion mutations found in the homology region in full length reads mapping to collapsed references. Data represent the mean of three biological replicates ± SEM with individual datapoints jittered. **D-E**, G-to-A mutation burden in proviral genomes from HeLa cells transduced with viral supernatants and harvested 72 hours later. D, Average number of G-to-A mutations in genomic strand 5’WGA motifs (cDNA strand 5’TCW) of CRV2.0 integrants. Data represent the mean of three biological replicates ± SEM with individual datapoints jittered. E, A3B mutation-burden separated into mean percentiles comparing A3B-mEGFP (red) to collapsing A3×3B-mEGFP (blue). **F-G**, A3B mutation burden in proviral genomes from 293T cells transduced with viral supernatants, treated with raltegravir, and harvested 24 hours later. F, Average number of G-to-A mutations in genomic strand 5’WGA motifs (cDNA strand 5’TCW) of CRV2.0 collapsed retroviruses. Data represent mean of three biological replicates ± SEM with individual datapoints jittered. G, A3B mutation-burden separated into mean percentiles comparing A3B-mEGFP (red) to collapsing A3×3B-mEGFP (blue).

Despite the improvement over non-CRV constructs, collapsing retrovirus integrants still showed an unacceptable G-to-A mutation burden due to A3B somehow being expressed in the producer cell. Long-read sequencing allows read-level grouping (*i.e*., genetic linkage) to ask if the A3B mutations cluster within a minority of reads or if they are relatively evenly distributed. By dividing reads into percentiles based on A3B mutagenic burden, this long-read approach showed that the majority of reads are unmutated, and the top 25% accounts for nearly the entire mutational burden (**Figure 3E**). The most likely explanation for this bimodal distribution is that the virus producing cells undergo VSV envelope-mediated autoinfection with newly produced CRVs, and that a subset of subsequently produced retroviruses are generated inadvertently in the presence of A3B expressed from this subset of nascently integrated constructs.

To test this producer cell self-infection hypothesis, viral stocks were produced in the presence of the retroviral integrase inhibitor raltegravir to prevent integration, and stocks were also collected at a much earlier timepoint (24 hours post-transfection versus 72 hours above). The average mutation burden in the non-collapsing A3B-mEGFP retrovirus remained high with 18.7 ± 1.5 APOBEC signature G-to-A mutations per viral genome (**Figure 3F**). In contrast, these technical modifications enabled a further reduction to 1.0 ± 0.05 APOBEC signature G-to-A mutations per viral genome in the collapsing CRV2.0 A3×3B-mEGFP condition (**Figure 3F**). Additionally, for CRV2.0, the APOBEC mutation burden is largely restricted to above the 96^th^ percentile (*i.e*., only 4% of viruses; **Figure 3G**). One possible explanation for this small fraction of residual hypermutated virus may be proviral DNA homologous recombination in the transfected virus-producing cell line shortly after transfection and before virus production.

### CRV-delivered A3B is nuclear and functional in target cells

Non-fluorescent CRVs were generated to validate the expression and functionality of untagged A3B constructs generated with this technology (illustrated in **Figure 4A**). Immunoblots and ssDNA deaminase activity assays demonstrate the expression and catalytic activity of non-fluorescent CRV1.1 and CRV2.1 constructs in cellular lysates (**Figure 4B, C**). Individual target cell A3B expression from CRV2.1 was also quantified by flow cytometry and shown to be nearly identical for catalytically active and inactive constructs (**Figure 4D**).

**Figure 4.**
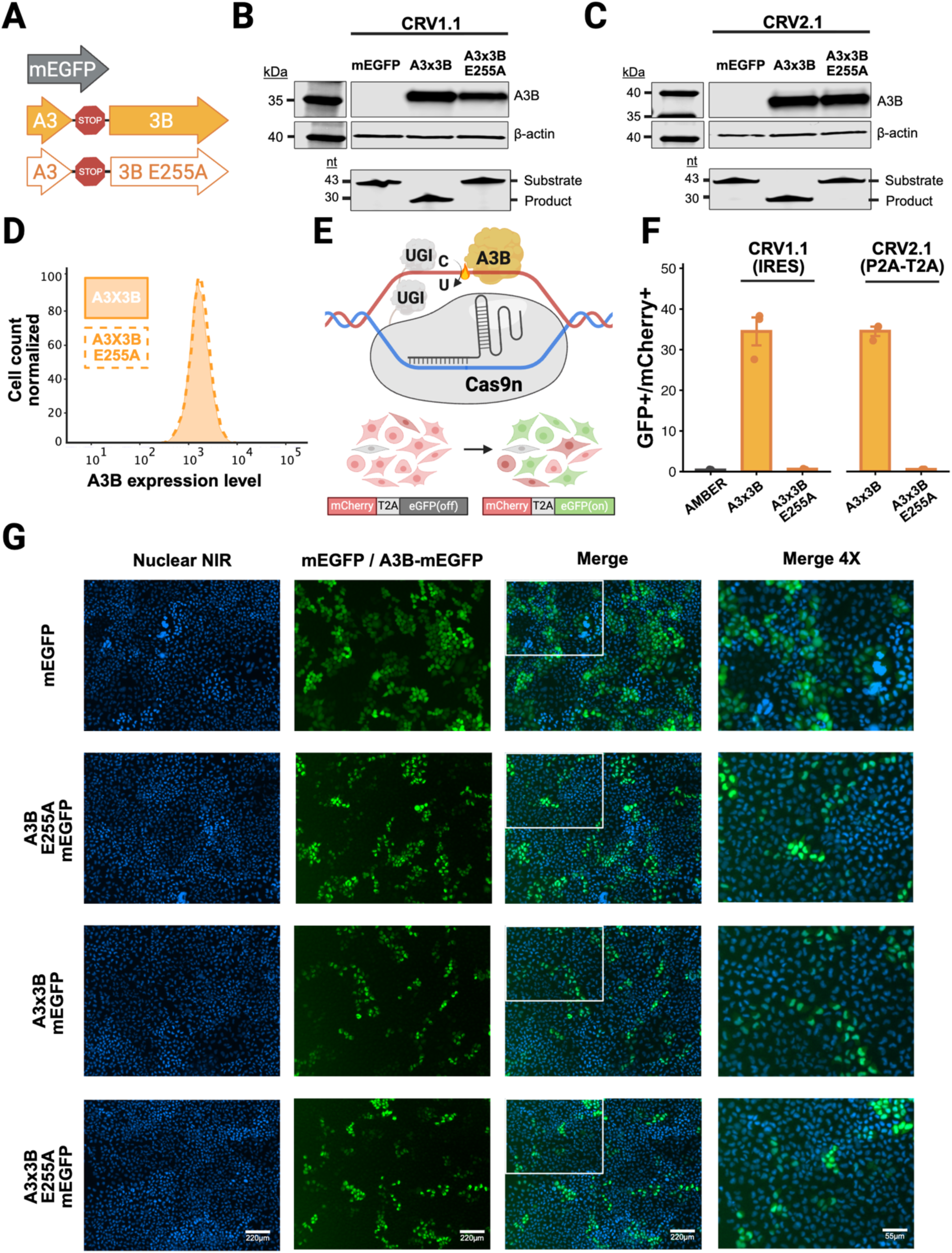
CRV-delivered A3B is nuclear and functional in target cells. **A**, Diagram of CRV1.0 and CRV2.0 constructs being tested here including mEGFP (gray), A3×3B (orange), and A3×3B-E255A (orange, white fill). **B-C**, Immunoblots and DNA deamination cleavage assay performed with 293T cell lysates following transduction with CRV1.0 or CRV2.0 constructs, respectively, illustrated in panel A. **D**, A3B expression levels in HeLa cells by flow cytometry >72 hours post-transduction with CRV2.0 A3×3B (solid orange) and A3×3B E255A (dashed orange). **E**, Illustration of AMBER, an episomal live-cell reporter for APOBEC3-catalyzed DNA deamination activity. Cas9 nickase/gRNA complex generates an R-loop in dual fluorescent reporter. A3B (orange) deamination followed by DNA damage repair of reconstitutes eGFP fluorescence. **F**, AMBER DNA editing quantification of 293T cells transduced with the indicated CRV1.0 and CRV2.0 constructs. Data represent the mean of three biological replicates ± SEM with individual datapoints overlayed and jittered. **G**, Live-cell imaging of HeLa cells transduced with standard retroviruses expressing mEGFP or A3B-E255A-mEGFP or in parallel with CRV1.0 constructs expressing A3B-mEGFP or A3B-E255A-mEGFP.

To assess the functionality of transduced A3B in living cells, we used our previously described APOBEC-mediated base editing reporter (AMBER) (Chen et al., 2026; Rieffer et al., 2023) (**Figure 4E**). Briefly, in this single cytosine base editing reporter system, a Cas9nickase(n)-Ugi-guide(g)RNA complex creates an R-loop spanning a mutant cytosine base in an eGFP cassette and then APOBEC-catalyzed deamination of this single target C-to-U (and nick-directed repair of the opposing DNA strand) results in a C-to-T mutation and restoration of eGFP fluorescence. Two copies of the uracil DNA glycosylase inhibitor protein (Ugi) are appended to the C-terminal end of the complex to prevent uracil base excision repair. An upstream mCherry reporter provides an internal control for the integrity of the reporter system. Importantly, C-to-U editing of the eGFP cassette in AMBER can be mediated from a deaminase tethered covalently to the Cas9n-gRNA complex (in *cis*) or from a deaminase expressed in *trans* following transfection or transduction (Chen *et al*., 2026) (a *trans* approach is depicted in **Figure 4E**). Here, active A3B expression in *trans* through transduction with CRV1.1 or CRV2.1 constructs results in similar frequencies of DNA C-to-U editing activity, as evidenced by 34.5 ± 1.2% and 33.5 ±3.4% eGFP/mCherry positive cells, respectively (**Figure 4F**). In comparison, catalytic mutant A3B-E255A constructs were unable to catalyze editing and restore fluorescence (**Figure 4F**). Consistent with strong DNA editing activity, parallel experiments with CRV2.0 constructs showed that A3B-mEGFP and A3B-e255A-mEGFP are localized to the nuclear compartment, whereas mEGFP alone is cell wide (**Figure 4F**).

## Discussion

Novel collapsing retroviruses (CRVs) were generated by splitting the A3B coding sequence into truncated tandem repeat elements with 1030 bp of homology which recombine in-frame at high efficiency to reconstitute a functional *A3B* minigene during reverse transcription. This technology effectively bypasses A3B-mediated virus restriction activity by eliminating functional protein expression in the virus producing cells. This approach leverages retroviral biology, which naturally requires two template-switching events to complete the reverse transcription process (*i.e*., first- and second-strand transfer reactions). The integrated *A3B* minigene is seamless, non-scarred, and nearly 100% error-free, and the protein product is fully functional and equivalent to the endogenous wildtype enzyme. Tethering drug selection to recombination removes sub-populations of cells expressing non-recombined and rarer frameshifted proviral integrants.

This work was inspired by earlier studies in which direct repeat system was used to circumvent restriction by A3G (Delviks-Frankenberry *et al*., 2019). In this system, a collapsing HIV-1 construct was used to deliver a Vif-resistant variant of A3G to a target cell population as a potential antiretroviral gene therapy strategy. Although the CRV system described here also leverages retroviral template switching to collapse direct repeats and remove a stuffer sequence, it is different in several fundamental ways. First, the CRV system is based on a MuLV backbone, not an HIV-1 backbone. Second, the CRV system uses human A3B as a test case because it is broadly relevant to virology as a virus restriction factor and to cancer biology as an endogenous mutagen. Third, CRV 2.0 and CRV2.1 utilize a C-terminal P2A-T2A tag to tether recombination to drug selection removing unwanted non-recombined viral integration events allowing for a homogenous transduced cell population. Fourth, the functionality of the CRV-delivered A3B enzyme is demonstrated in a target cell population through editing of a single DNA target cytosine in an eGFP biosensor.

The ability to efficiently introduce viro-toxic wildtype *A3B* transgenes and mutant derivatives into cellular systems has broad utility by enabling exogenous A3B expression studies, complementation experiments, and variant analyses. It also enables, for instance, delivery of an entire deaminase-Cas9n-Ugi-gRNA base editing complex to a target cell population for purposes of precision genome engineering for research and therapeutic purposes. Alternative existing approaches to stably introduce *A3B* minigenes suffer from inherent limitations. Stable transfection and transposase-based systems are hindered by low efficiency particularly in difficult to transfect cell lines (Burns et al., 2013; McCann et al., 2023; Stenglein and Harris, 2006). Alternative approaches utilize retroviruses but seek to limit A3B expression in other ways. One approach relied on transgenes that express *A3B* under Tet/Dox-repressible promotors such that virus can be generated in Tet/Dox-expressing cell lines to limit *A3B* expression (Buisson et al., 2017; Ortega et al., 2024; Ortega et al., 2025). This approach may suffer from leaky A3B expression, because Tet/Dox repression is typically overwhelmed by the high numbers of plasmids per cell generated by transient transfection. Another approach is to introduce a minigene flanked by *loxP* sites such that *A3B* expression is Cre recombinase-inducible (Garcia et al., 2025; Horisawa et al., 2025). A drawback here is the additional requirement for Cre recombinase expression, the efficiency of which may introduce further unwanted variability. Finally, A3B retrovirus restriction can be overcome by concentrating virus stocks to try to enhance viral titers [*e.g*., (Carpenter *et al*., 2023; Huff et al., 2018; Zhang et al., 2026)]. However, as shown here with long-read sequencing, this brute force method is limited by the introduction a large percentage of low fidelity, mutated transgenes that can complicate experiments and mask the true impact of A3B functionality.

In our studies with CRV technology here, leaky *A3B* expression due to autoinfection of the retrovirus producing cell population led to a subset of self-hypermutated *A3B* transgenes in the target cell population. This was mostly mitigated by generating viral stocks in the presence of antiretroviral drugs like the integrase inhibitor raltegravir and by collecting viral supernatants at early timepoints prior to autoinfection. This risk can be further mitigated by utilizing a combination of antiretroviral drugs, optimized retrovirus producing cell lines that lack the endogenous *APOBEC3* locus, and producing cell lines engineered to lack the LDL-R utilized for the entry of VSV-G envelope-pseudotyped particles. Although A3B was used as an extreme test case here (extremely toxic to retroviruses), the CRV system may have broader utility for delivering other viro-toxic factors such as other restriction factors and base editing complexes into desired target cell populations.

### Limitations of the study

The current CRV1.0 and 2.0 constructs enable viro-toxic cargo to be delivered efficiently to a target cell population, but their constitutive expression may cause target cell cytotoxicity (especially over long-term experiments). This could be alleviated in future constructs by coupling the CRV technology to an inducible system and/or to a degron. The current CRV systems are MLV-based and therefore only amendable to transgene delivery to dividing cells. HIV (lenti)-based CRV’s could alleviate this additional limitation and further expand utility in non-dividing cell types such as neurons.

## Resource availability Lead contact

Further information and requests for resources should be directed to and will be fulfilled by the lead contact, Reuben Harris.

## Materials availability

The CRV constructs reported here are available from Addgene (catalog numbers available after acceptance) and/or by email request to the lead contact.

## Data and code availability

Long-read sequencing data are available upon request to the lead contact. Original code for long-read sequencing alignment and analysis can be found on GitHub at: https://github.com/mullallyc/Collapsing-retroviruses-for-efficient-delivery-of-viro-toxic-cargoe

Any additional information required to reanalyze the data reported in this work paper is available from the lead contact upon email request.

## Acknowledgments

We thank members of the Harris laboratory for support and constructive feedback. BioRender was used to create the Graphical Abstract, Fig. 1A, Fig. 1B, Fig. 2A, Fig. 2B, Fig. 3A, and Fig. 4A. Fig. 4E, and Fig. S1.

## Author contributions

Conceptualization, C.D.M., R.S.H.; Methodology, C.D.M., M.A.C.; Formal analysis, C.D.M., B.S., Y.C., H.B.G., M.A.C., R.S.H.; Investigation, C.D.M., B.S., Y.C., H.B.G., M.A.C., R.S.H.; writing – original draft, C.D.M., R.S.H. writing – review & editing, all authors; resources, C.D.M., R.S.H.; funding acquisition, C.D.M., R.S.H.; supervision, R.S.H.

## STAR★Methods

Detailed methods are provided in the online version of this paper and include the following:

- KEY RESOURCES TABLE
- EXPERIMENTAL MODEL AND STUDY PARTICIPANT DETAILS
- METHOD DETAILS
- QUANTIFICATION AND STATISTICAL ANALYSIS

