## Supplementary material for "Collapsing retroviruses for efficient delivery of viro-toxic cargoes": SI Figs S1-S3

**Supplementary Figure 1.** Schematic of reverse transcription and second strand synthesis of CRV1.0

**Supplementary Figure 2.** Data from HeLa cells transduced with CRV2.0

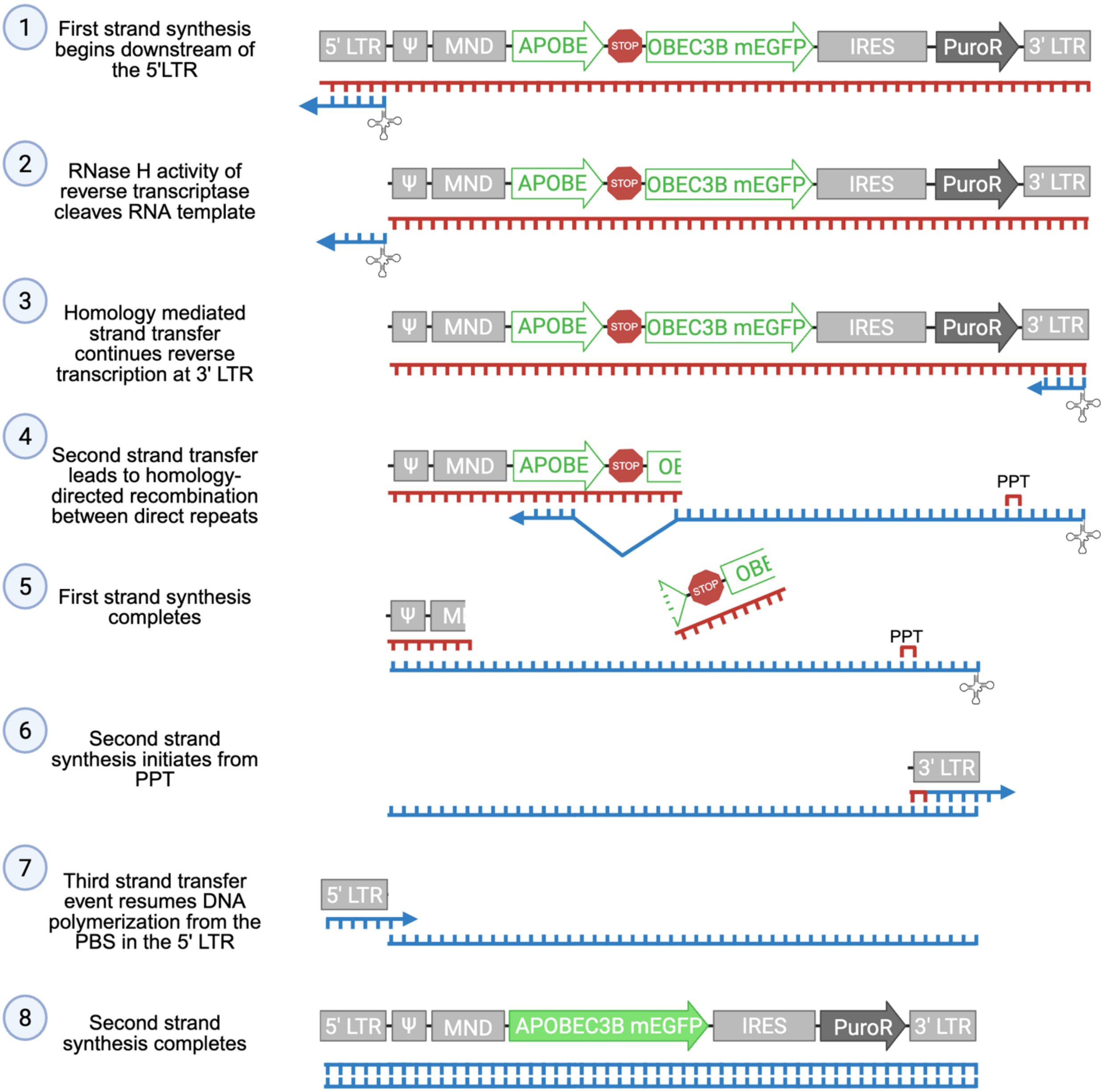
1
First strand synthesis begins downstream of the 5'LTR
5' LTR
Ψ
MND
APOBE
STOP
OBEC3B mEGFP
IRES
PuroR
3' LTR
2
RNase H activity of reverse transcriptase cleaves RNA template
3
Homology mediated strand transfer continues reverse transcription at 3' LTR
4
Second strand transfer leads to homology-directed recombination between direct repeats
PPT
5
First strand synthesis completes
6
Second strand synthesis initiates from PPT
7
Third strand transfer event resumes DNA polymerization from the PBS in the 5' LTR
8
Second strand synthesis completes
APOBEC3B mEGFP

**Supplementary Figure 1. Schematic of reverse transcription and second strand synthesis of CRV1.0**

RNA viral genome (red) is reverse transcribed into proviral DNA following the steps indicated. Both first- and second-strand synthesis require template switching events. CRV collapse is mediated by an additional template switch during first-strand synthesis. CRV2.0 constructs function similarly except P2A-T2A replaces the IRES.

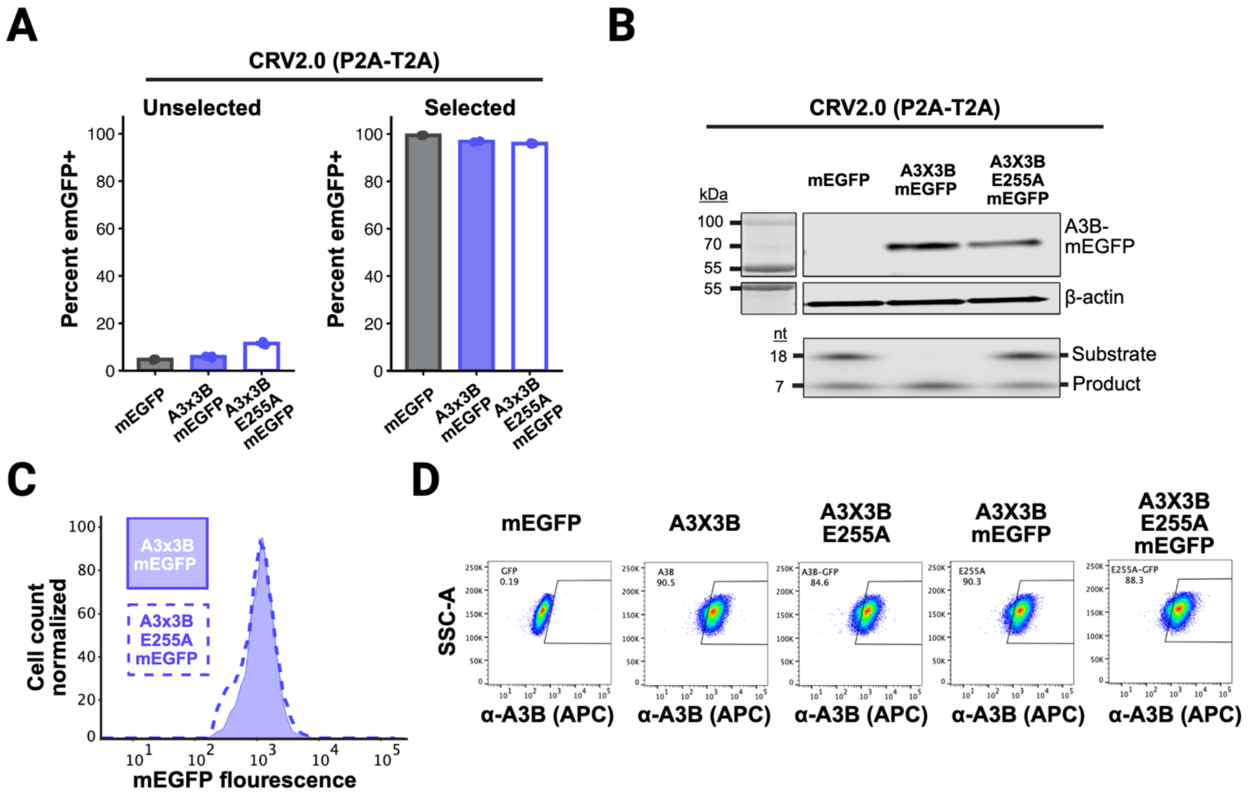
A
CRV2.0 (P2A-T2A)
Unselected
Selected
Percent emGFP+
mEGFP
A3x3B mEGFP
A3x3B E255A mEGFP
B
CRV2.0 (P2A-T2A)
kDa
100
70
55
55
mEGFP
A3X3B mEGFP
A3X3B E255A mEGFP
A3B-mEGFP
β-actin
nt
18
7
Substrate
Product
C
Cell count normalized
A3x3B mEGFP
A3x3B E255A mEGFP
mEGFP flourescence
D
mEGFP
A3X3B
A3X3B E255A
A3X3B mEGFP
A3X3B E255A mEGFP
SSC-A
α-A3B (APC)

**Supplementary Figure 2. Data from HeLa cells transduced with CRV2.0**

**A**, Quantification of percent mEGFP+ of HeLa cells transduced with CRV2.0 mEGFP, A3x3B-mEGFP, or A3x3B-E255A-mEGFP pre- and post-selection. Data represent the mean of three biological replicates ± SEM with individual datapoints jittered.

**B**, Immunoblots and DNA deamination cleavage assay results using HeLa cell lysates following transduction with the indicated CRV2.0 constructs (post-selection). Levels of A3B activity in CRV2.0 A3B-expressing cells are sufficient to promote full deamination of the 18nt oligo substrate, whereas the lower, residual deamination activity of the control reactions is likely due to endogenous A3B.

**C**, A3B- and A3B-E255A-mEGFP expression levels in live HeLa cells by flow cytometry [CRV2.0 A3x3B (blue) and A3x3B E255A (dashed blue)].

**D**, Flow cytometry dot plots of HeLa cells transduced with the indicated CRV2.0 and CRV2.1 constructs and stained with a specific anti-A3B monoclonal antibody.
