## Supplementary material for "Collapsing retroviruses for efficient delivery of viro-toxic cargoes": Methods

### STAR★Methods

### KEY RESOURCES TABLE

| REAGENT or RESOURCE | SOURCE | IDENTIFIER |
|---|---|---|
| **Antibodies** | | |
| Rabbit anti-A3B monoclonal antibody | Cell Signaling Technology | Cat# 41494 |
| Anti-human Fcg receptor blocker | BioLegend | Cat# 422302 |
| Anti-rabbit Alexa Fluor 647-conjugated antibody | Thermo Fisher Scientific | Cat# A32733 |
| Anti-actin antibody | Sigma-Aldrich | Cat# A1978 |
| Goat anti-rabbit HRP conjugated antibody | Cell Signaling Technology | Cat# 7074 |
| Goat anti-mouse IRDye 680LT antibody | LI-COR | Cat# 926-68020 |
| Bacterial Strains | | |
| NEB Stable Competent *E. coli* | New England Biolabs | Cat# C304 |
| Critical Commercial Assays | | |
| Quick-RNA Viral Kit | Zymo Research | Cat# R1035 |
| QuickExtract™ DNA Extraction Solution | LGC Biosearch Technologies Products | Cat# LGCQE09050 |
| MycoAlert Mycoplasma Detection Kit | Lonza | Cat# LT07 |
| ZymoScript RT PreMix Kit | Zymo Research | Cat# R3012 |
| SsoFast™ EvaGreen Supermix | Bio-Rad | Cat# 1725200 |
| Monarch® Spin PCR & DNA Cleanup Kit | New England Biolabs | Cat# T1130 |
| Bradford assay | Sigma-Aldrich | Cat# B6916 |
| Experimental models Cell lines | | |
| 293T | ATCC | Cat# CRL-3216 |
| Expi-293F | Gibco | Cat# A14527 |
| HeLa | ATCC | Cat# CCL-2 |

| Chemicals, peptides, and recombinant proteins | | |
|---|---|---|
| AgeI-HF | New England Biolabs | Cat# R3552 |
| T4 DNA ligase | New England Biolabs | Cat# M0202 |
| NotI | New England Biolabs | Cat# R0189 |
| BspEI | New England Biolabs | Cat# R0540 |
| PEI MAX | Thermo Scientific | Cat# 047336.03 |
| Raltegravir | MedChem Express | Cat# HY-10353 |
| Puromycin dihydrocloride | Gibco | Cat# A1113803 |
| TransIT LT-1 | Mirus Bio | Cat# MIR 2304 |
| UDG | New England Biolabs | Cat# M0280 |
| repliQa HIFI | Quantabio | Cat# 95200 |
| RNase A | New England Biolabs | Cat# M0314 |
| cOmplete protease inhibitor | Roche | Cat# 11697498001 |
| **Plasmid constructs** | | |
| AMBER Reporter Plasmid | Addgene | Cat # Pending |
| Cas9n-BE4max plasmid | Addgene | Cat # Pending |
| AMBER gRNA plasmid | Addgene | Cat # Pending |
| muLV-MND-A3x3B-IRES-PuroR | This study; Addgene | Cat # Pending |
| muLV-MND-A3x3B-E255A-IRES-PuroR | This study; Addgene | Cat # Pending |
| muLV-MND-A3x3B-P2A-T2A-PuroR | This study; Addgene | Cat # Pending |
| muLV-MND-A3x3B-E255A-P2A-T2A-PuroR | This study; Addgene | Cat # Pending |
| muLV-MND-A3x3B-mEGFP-IRES-PuroR | This study; Addgene | Cat # Pending |
| muLV-MND-A3x3B-E255A-mEGFP-IRES-PuroR | This study; Addgene | Cat # Pending |
| muLV-MND-A3x3B-mEGFP-P2A-T2A-PuroR | This study; Addgene | Cat # Pending |

| | | |
|---|---|---|
| muLV-MND-A3x3B-E255A-mEGFP-P2A-T2A-PuroR | This study; Addgene | Cat # Pending |
| muLV-MND-mEGFP-P2A-T2A-PuroR | This study; Addgene | Cat # Pending |
| muLV-MND-mEGFP-IRES-PuroR | This study; Addgene | Cat # Pending |
| **Software and algorithms** | | |
| FlowJo 10.10.1 | BD Biosciences | RRID:SCR_008520; https://www.flowjo.com |
| ChatGPT Codex | OpenAI | https://chat.openai.com |
| Minimap2 v2.30 | Heng Li | RRID:SCR_018550; https://github.com/lh3/minimap2 |
| Pysam v0.23.3 | Pysam developers | RRID:SCR_021017; https://github.com/pysam-developers/pysam |
| Python V3.12 | Python Software Foundation | RRID:SCR_008394; https://www.python.org |
| R v4.5.0 | R Foundation for Statistical Computing | RRID:SCR_001905; https://www.r-project.org |
| Custom Analysis Scripts | This Paper | https://github.com/mullallyc/Collapsing-retroviruses-for-efficient-delivery-of-viro-toxic-cargoe |
| **Oligonucleotides** | | |
| Direct repeat A3 (A3B) fwd | This study | actgacgcggccgcatgaatccacagatcagaaatccgatggag |
| Direct repeat A3 (A3B) rev | This study | gcgtggtactgaggcttgaacaattgtcattaaccggttcaccagggctggaagggacatc |

| | | |
|---|---|---|
| Direct repeat 3B (A3B) fwd | This study | gatgtcccttccagccctggtgaaccggttaatgacaa ttgttcaagcctcagtaccacgc |
| Direct repeat 3B (A3B) rev | This study | gtacactccggattagtttccctgattctggagaatgg |
| Collapse A3X3B repeats fwd | This study | gatgtcccttccagccctgggatggactagag |
| Collapse A3X3B repeats rev | This study | ggaagggacatccctggcggtacacaaagg |
| Delete A3x3B from A3x3B-GFP Fwd | This study | gccgcatggtgagcaagggcgaggagctgttc |
| Delete A3x3B from A3x3B-GFP Rev | This study | tcaccatgcggccgcggatcaattcctgc |
| No raltegravir amplicon primer fwd | This study | cagccctcactccttctctagg |
| No raltegravir amplicon primer rev | This study | tcaggcaccgggcttgcg |
| Raltegravir amplicon primer fwd | This study | tcgatcctccctttatccagccctcactcc |
| Raltegravir amplicon primer rev | This study | gagccatggggtcgtgcgctcctttc |
| Deaminase assay linear substrate | This study | ATTATTATTATTCTAATGGATTTA TTTATTTATTTATTTATTT-/36-FAM/ |
| Deaminase assay hairpin substrate | This study | TGCCATCTATCGATGGGA/36-FAM/ |
| **Other** | | |
| Amicon Ultra Centrifugal Filter, 30 kDa MWCO | Millipore | Cat# UFC903008 |
| Opti-MEM | Gibco | Cat# 31985062 |
| RPMI-1940 medium | Thermo Fisher Scientific | Cat# 11875093 |
| Expi293™ Expression Medium | Gibco | Cat# A1435102 |
| Fetal Bovine Serum | Biowest | Cat# 058N24 |
| DMEM | Thermo Fisher Scientific | Cat#11-965-118 |
| 384 well plate | Genesee Scientific | Cat# 240308W |

| | | |
|---|---|---|
| Incucyte® Nuclight NIR Lentivirus | Sartorius | Cat# 4805 |
| Polyvinylidene difluoride (PVDF) Immobilon-FL membrane | Millipore | Cat# IPFL00005 |
| Casein blocking buffer | Sigma-Aldrich | Cat# B6429 |
| Ampure XP Beads | Beckman Coulter | Cat# A63881 |

### EXPERIMENTAL MODEL AND STUDY PARTICIPANT DETAILS

The following human cancer cell lines were used: 293T cells (ATCC, Cat#: CRL-3216), Expi-293F cells (Gibco, Cat#: A14527), and HeLa cells (ATCC, Cat#: CCL-2). There was no need for study participants.

### METHOD DETAILS

#### Plasmids

The parental MLV constructs used to generate A3x3B CRV1.0 have been described (Carpenter *et al.*, 2023). The tandem-repeat A3x3B was generated by PCR amplification from previously reported expression plasmids (Auerbach et al., 2022) containing a wildtype *A3B* open reading frame or a catalytically mutant A3B-E255A, with each ORF disrupted by an intron positioned between exons 5 and 6 (see Table 1 for primer sequences). This amplification produced N- and C-terminal open reading frame fragments corresponding to A3B residues 1-359 and 61-382, as well as the intron, totaling 1030 bp of self-homology. In addition, these fragments contained an appended 3× stop-codon stuffer sequence. The PCR fragments were digested with AgeI-HF (NEB, Cat#R3552) and ligated with T4 DNA ligase (NEB, Cat#M0202). The ligated insert was subsequently cloned into the base vector by standard restriction-ligation cloning using NotI (NEB, Cat#R0189) and BspEI (NEB, Cat#R0540). Ligated constructs were transformed into NEB Stable Competent *E. coli* (NEB, Cat#:C3040). NEB stable cells were used for all subsequent plasmid recovery and expansion. C-terminal mEGFP fusion tags were synthesized and cloned by Genscript. Subsequent CRV2.0 derivatives of this construct containing a P2A-T2A C-terminal tag rather than an IRES were synthesized and cloned by GenScript. Vector and sequence matched non-collapsing A3B control as well as mEGFP alone control plasmids were generated by site directed mutagenesis intended to collapse vectors (see Table 1 for primer sequences). All plasmid constructs are including maps are available on Addgene (see below).

**Cell culture**

293T cells (ATCC, Cat#: CRL-3216) were maintained in RPMI-1940 medium (Thermo Fischer Scientific, Cat#: 11875093) supplemented with 10% (v/v) Fetal Bovine Serum (Biowest, Cat#: 058N24) at 37C with 5% $CO_2$. Expi-293F cells (Gibco, Cat#: A14527) were maintained in Expi293™ Expression Medium (Gibco, Cat#: A1435102) at 37 $^{o}$C with 8% $CO_2$ shaking at 125 rpm. HeLa cells (ATCC, Cat#: CCL-2) were maintained in DMEM (Thermo Fischer Scientific, Cat#:11-965-118) supplemented with 10% (v/v) Fetal Bovine Serum at 37$^{o}$C with 5% $CO_2$. Cultures were routinely verified as mycoplasma-negative by screening with the MycoAlert Mycoplasma Detection Kit (Lonza, Cat#: LT07).

**Virus production**

Expi293-F suspension cells were seeded at $1.5 \times 10^6$ cells/mL in 30 mL of medium. Twenty-four hours later, the cells were transfected with the viral production plasmids: 5 µg of VSV-G, 10 µg of MLV gag-pol, and 15 µg of the collapsing or non-collapsing retroviral vector, for a total of 30 µg of DNA. The plasmid DNA was diluted in 3 mL of Opti-MEM (Gibco, Cat#: 31985062), 160 µL of PEI MAX (40 kDa, 1 mg/mL; Thermo Scientific, Cat#: 047336.03) was added, and the mixture was mixed gently by pipetting. The transfection mix was incubated for 10 min and then added dropwise to the Expi293-F cells. Viral supernatant was harvested at 72 hours and filtered in a 45µm filter. Viral stock was concentrated using Amicon Ultra Centrifugal Filter, 30 kDa MWCO (Millipore, Cat#: UFC903008), spinning at 4,500 x *g* for 15 minutes followed by one 10 mL PBS wash. Viral stocks were then resuspended in PBS and flash frozen in 1.5mL tubes using liquid nitrogen before long-term storage at -80$^{o}$C.

For subsequent virus production, adherent 293T cells were seeded in T75 flasks at a density of $2.1 \times 10^6$ cells in full-growth media containing 1µM raltegravir (MedChemExpress, Cat#: HY-10353). Cells were transfected with viral production plasmids after 24 hours. Viral production plasmids include 1.66 µg of VSV-G containing, 3.33 µg of mLV gag-pol containing and 5µg of non-collapsing or collapsing retroviral vectors. Plasmids were diluted in 1.5mL of Opti-MEM. 3µl of Transit LT-1 (Mirus Bio, Cat#: MIR 2304) was added per 1µg of DNA, followed by gentle mixing by pipetting. After a 15 minute incubation, transfection mix was added dropwise to the 293T.

Viral supernatant was harvested after 24 hours and filtered with a 45µm filter. Amicon Ultra Centrifugal Filter, 30 kDa MWCO, spinning at 4,500 x *g* for 15 minutes followed by three 10mL PBS washes. Viral stocks were then resuspended in residual PBS and flash frozen in 1.5mL tubes using liquid nitrogen before long-term storage at -80°C.

**Viral stock quantification**

Viral stocks were divided into three technical replicates, and RNA was isolated using Quick-RNA™ Viral Kit (Zymo Research, Cat#: R1035) with a final elution volume of 40uls. 1uls was carried forward for reverse transcription was performed with ZymoScript RT PreMix Kit (Zymo Research, Cat# R3012), final volume of 10uls. The resultant cDNA mix was diluted 1:10 and 1uls was added to qPCR mix containing SsoFast™ EvaGreen Supermix (Bio-Rad, Cat#: 1725200) and the primer set (5'TCGTAGAAGGGGAGGTTGC3', 5'ACCAGGGCAAGGGTCTG3') at 300nM final concentration in 5 µl final volume. qPCR was run in a 384 well plate (Genesee Scientific, Cat#: 240308W) and run on a LightCycler 480 II (Roche, Cat#: 05015243001). No-RT controls were run in parallel to account for residual trace DNA amplification. Each sample was divided into 4 technical replicates, averaged and the viral concentration was determined using a standard curve measured in parallel. DNA input for standard curve was linear DNA product generated by PCR amplification of 100pg collapsing retroviruses plasmid template with repliQa HIFI ToughMix (Quantabio, Cat#: 95200) and the same primers at a final concentration of 300nM, final volume 50uls. DNA product was cleaned up with Monarch® Spin PCR & DNA Cleanup Kit (New England Biolabs, Cat#:T1130) and quantified with the NanoDrop™ One spectrophotometer (ND-ONE-W). Standard curve was measured in parallel by $\log_{10}$ serial dilution of PCR starting at 10pg of input.

**Live-cell virus titration experiments**

HeLa cells were first transduced with Incucyte® Nuclight NIR Lentivirus (EF-1α, Puro; Sartorius, Cat#: 4805) according to the manufacturer's recommendations, and transduced cells (HeLa-NIR) were selected with puromycin at 2 μg/mL. This construct drives nuclear-restricted expression of the near-infrared protein iRFP713, which provides a non-perturbing nuclear label that enables automated cell counting during live-cell imaging. HeLa-NIR cells were seeded into 96-well plates at 9,375 cells/cm². The following day, cells were infected in triplicate with a nine-point, two-fold serial dilution series of collapsed A3B-mEGFP or A3B-E255A-mEGFP or non-collapsed A3x3B-mEGFP or A3x3B-E255A-mEGFP virus in the presence of 4 μg/mL polybrene, with mEGFP fluorescence serving as the readout for infection. Plates were immediately transferred to an Incucyte® live-cell analysis system, and the NIR (nuclei) and green (mEGFP) channels were imaged every 3 hours with 10X objective. Approximately 24 h after infection, the inoculum was removed and replaced with fresh medium, and imaging was continued for a total of 72 h. Infection was quantified as the number of mEGFP-positive cells by live-cell flow cytometry (see below).

**Recombination rate assay**

293T and HeLa cells were seeded in 12-well plates at 125,000 cells per well. After 24 hours, they were transduced at low MOI (<0.1) with the 72 hour viral prep made from each of the indicated constructs. After 36 hours, the cells were split 1:2 and one unselected and one selected at 2ug/mL puromycin dihydrocloride (Gibco, Cat#: A1113803) for at least 72 hour and until all mock transduced cells were completely selected and non-viable. Representative images were taken (see below) before cells were trypsinized with 0.05% Trypsin EDTA (Gibco, Cat#: 25300-054) and resuspended in complete growth media and per cell GFP fluorescence was measured by flow cytometry (see below).

**Live-cell imaging**

Representative live cell images were acquired using an Incucyte SX5 Live-Cell Analysis System housed in a humidified tissue culture incubated at 36.5C and 5% $CO_2$. Images were collected in brightfield, green fluorescence and near infrared fluorescence channels using either

10X or 20X objectives. Image acquisition settings were held constant across experiments. Images shown are representative of three independent biological replicates.

**Flow cytometry**

Flow cytometry was used as a single-cell resolution quantitative measurement of expression of A3B-mEGFP and the catalytically inactive A3B-E255A-mEGFP expressed from standard non-collapsing, CRV1.0 and CRV2.0 vectors, using mEGFP fluorescence as a surrogate for construct expression. Cells had undergone full drug selection prior to the experiment. Cells were washed once in PBS, detached by trypsinization, and resuspended as a single-cell suspension in PBS prior to acquisition. Samples were run on a LSRFortessa X-20 (BD Biosciences) flow cytometer, with a minimum of 10,000 events recorded per sample. A hierarchical gating strategy was applied: cells were first gated on FSC-A versus SSC-A to exclude debris and define the main cell population, and single cells were then selected by sequential doublet exclusion on SSC-W versus SSC-H followed by FSC-W versus FSC-H. mEGFP signal was resolved on histogram of GFP-A and the mEGFP+ population was defined as signal above non-fluorescent negative control cells. A3B expression was compared across constructs based on the distribution of mEGFP fluorescence intensity (MFI) of this population. Live cell viral titer and recombination rate was determined as the percentage cells mEGFP+. Data were analyzed in softwares FACSDiva and FlowJo 10.10.1 (BD Biosciences).

A3B expression from untagged and mEGFP tagged vectors was independently measured by flowcytometry after fixation and immunostaining. HeLa cells were transduced with CRV2.0 and 2.1 vectors expressing mEGFP, A3B, A3B E255A, A3-mEGFP, or A3B E255A mEGFP cells and selected with 2ug/mL puromycin dihydrocloride for at least 72 hours. Wells were trypsinized and seeded at 0.4 x $10^6$ cells per well in a sterile 6-well plate and cultured under identical conditions for 48 hours. The cells were then harvested by trypsinization, and 1 x $10^6$ cells were processed for subsequent analyses. Cells were washed once with PBS and incubated with 200 ml of 1:200 anti-human Fcg receptor blocker (BioLegend, Cat#: 422302) diluted in flow buffer (2% FBS in PBS) for 30 min at 4°C. Following two PBS washes, cells were permeabilized and fixed by dropwise addition of 400 ml ice-cold methanol, while vortexing, and incubated at 4°C for 20 minutes. Cells were subsequently washed twice with PBS to remove methanol and for rehydration.

Cells were then blocked with 1% BSA in PBS for 10 minutes at room temperature and then incubated overnight at 4°C with either blocking buffer alone or with 1 mg anti-A3B specific antibody (Cell Signaling Technology, Cat#: 41494) diluted in blocking buffer. The next day, cells were washed 3X with PBS and incubated with 1:1000 anti-rabbit Alexa Fluor 647-conjugated secondary antibody (Thermo Fisher Scientific, Cat#: A32733) for one hour at 4°C, diluted in blocking buffer. After washing, cells were fixed in 0.5% formalin and data acquired and analyzed using LSRFortessa X-20 (BD Biosciences 656385) using FACSDiva and FlowJo 10.10.0 software (BD Biosciences).

**Immunoblots**

Cells had undergone full drug selection prior to the experiment. For every 1 million cells, the pellet was resuspended in 100 µL of HED buffer (25 mM HEPES, 5 mM EDTA, 10% glycerol, 1 mM DTT, and 1× cOmplete protease inhibitor; Roche, Cat#: 11697498001). Cells were lysed by snap-freezing followed by a 20-min sonication in a water-bath sonicator, and the lysates were clarified by centrifugation at 16,000 × *g* for 15 min. Total protein was quantified in triplicate by Bradford assay (Sigma-Aldrich, Cat#: B6916), with each sample diluted 1:100 into the assay reagent, and absorbance was read at 595 nm on a Tecan Spark multimode plate reader. Samples were mixed with SDS-PAGE loading buffer (125 mM Tris, pH 6.8, 5% SDS, 30% glycerol, 100 mM dithiothreitol [DTT], and Orange G dye), normalized to equal concentrations, and denatured at 98 °C for 10 min prior to immunoblotting. Proteins were resolved on a 4–20% gradient SDS-PAGE gel at 130 V for 90 min and then transferred onto a polyvinylidene difluoride (PVDF) Immobilon-FL membrane (Millipore, Cat#: IPFL00005) using a Bio-Rad rapid-transfer apparatus according to the manufacturer's instructions. Membranes were blocked in casein blocking buffer (Sigma-Aldrich, Cat#: B6429) for 1 h to prevent nonspecific binding, then probed overnight at 4 °C with primary antibody. The primary antibodies were rabbit anti-human A3B (1:1000; Cell Signaling Technology, Cat#: 41494) and anti-actin (1:5000; Sigma-Aldrich, Cat#: A1978). After primary incubation, membranes were washed three times in PBST (10 min each) and incubated for 1 h with the appropriate secondary antibody, goat anti-rabbit HRP (1:5000; Cell Signaling Technology, Cat#: 7074) or goat anti-mouse IRDye 680LT (1:5000; LI-COR, Cat#: 926-68020). Membranes were then washed a further three times in

PBST and imaged on a LI-COR Odyssey M imager.

**DNA deamination assays**

Single-stranded DNA cytidine deaminase activity in whole-cell lysates was measured using a fluorescence-based oligonucleotide cleavage assay. 293T and HeLa cells were lysed in HED buffer (25 mM HEPES, 5 mM EDTA, 10% glycerol, 1 mM DTT, and 1× cOmplete protease inhibitor), and 7.5µg of cleared lysate was used per reaction. Each 20 µL reaction contained 4 pmol of a 3′-FAM–labeled single-stranded DNA substrate, 0.025 U uracil-DNA glycosylase (UDG; NEB, Cat# M0280), and 1.75 U RNase A (NEB, Cat#: M0314). The substrate was matched to the virus being assayed: for the untagged virus, 5′-ATTATTATTATTCTAATGGATTTATTTATTTATTTATTTATTT-FAM, incubated at 37 °C for 2 h; for the mEGFP-tagged virus, 5′-TGCCATCTATCGATGGGA/36-FAM/, incubated at 37 °C for 4 h. Reactions were then treated with 100 mM NaOH and heated at 95 °C for 10 min to cleave the backbone at abasic sites. Products were resolved on 15% denaturing TBE–urea polyacrylamide gels, run at 12 V for 45 min, and FAM fluorescence was imaged on a LI-COR Odyssey M.

**AMBER assays**

Untethered AMBER assays were performed as described (Chen *et al.*, 2026). 293T cells were seeded in 12-well plates at 125,000 cells per well. After 24 hours, they were transduced at low MOI (<0.1). After 36 hours, the cells were selected at 2ug/mL puromycin for at least 72 hour and until all mock transduced cells were completely selected and non-viable. Cells were trypsinized with 0.05% Trypsin EDTA and replated at 125,000 cells per well in 12-well plates and were transfected 24 hours plated. Transfection mix contained 300 ng of AMBER reporter plasmid, 200 ng of Cas9n-BE4max plasmid, 100 ng of AMBER gRNA plasmid and 400 ng of empty vector pcDNA3.1 diluted in 100uls of optiMEM. TransIT-LT1 was added at a ratio of 3uls per ug of DNA followed by gentle pipette mixing and a 15-minute incubation. Transfection mix was added dropwise to the cells. At 48 h, cells were trypsinized and suspended in 1 ml of complete growth media. An aliquot of 200 µl was used for flow cytometry to quantify mCherry and eGFP expression with a minimum of 10,000 events recorded per sample A hierarchical gating strategy was applied: cells were first gated on FSC-A versus SSC-A to exclude debris and define the main cell population, and single cells were then selected by sequential doublet exclusion on SSC-W versus SSC-H followed by FSC-W versus FSC-H. mCherry signal was resolved on histogram of mCherry-A and the mCherry+ population was defined as signal above non-fluorescent negative control cells. Subgating on mCherry+ population, GFP+ was determined in the same manner. AMBER activation is reported as (GFP+ and mCherry+)/mCherry+.

**Long read sequencing**

HeLa and 293T cells were seeded at 10,000 cells per well in a 96 well plate and transduced after 24 hours. DNA was harvested after 72 hours using QuickExtract$^{TM}$ DNA Extraction Solution (LGC Biosearch Technologies Products, Cat#: LGCQE09050) following their protocol. 0.5ul of extracted genomic DNA was PCR amplified with repliQa HIFI ToughMix at a final concentration of 300nM, final volume 20uls (see primers in Table 1). PCR product was purified using adding 0.8X Ampure XP Beads (Beckman Coulter, Cat#: A63881). Long read sequencing libraries were prepared and sequenced by Plasmidsaurus with v14 library prep chemistry enabling minimal amplicon fragmentation. FASTQ files were analyzed using a custom Python pipeline. Portions of the custom Python analysis code were drafted with assistance from ChatGPT 5.5 (OpenAI). All code was reviewed, modified, and validated by the authors before use. Reads alignment was performed with minimap2 (Li, 2018) using the Oxford Nanopore preset (-x map-ont) with SAM output (-a). SAM alignments were parsed with pysam (Li et al., 2009) to extract alignment metrics, base-level read-to-reference mappings, CIGAR information, and per-base quality scores for mutation and INDEL calling. Reads were aligned to corresponding reference maps for either collapsed or non-collapsed vectors. Only primary alignments and near full-length alignments to the expected amplicon were included in the final analyses. Reads were retained if the met all the following criteria: alignment identity ≥80%, aligned query length/reference length ≥90%, reference span/reference length ≥90%, and aligned query length/read length ≥90%. These thresholds were intended to retain full-length reads while allowing for the inclusion of highly mutated reads.

For recombination-rate analysis, reads were competitively and exclusively aligned to collapsed and non-collapsed reference maps. Unmapped and ambiguous reads were excluded from the recombination denominator, and recombination rate was calculated as collapsed reads/ (collapsed reads + non-collapsed reads).

For INDEL analyses, reads which exclusively mapped to the collapsed reference using the same final alignment filters were considered. To reduce sequencing-error artifacts, only high-confidence INDELs were counted. For deletions, the flanking 5′ and 3′ reference-adjacent read bases were required to have Phred quality >30. For insertions, both flanking bases were required to have Phred quality >30, and the mean Phred quality of the inserted bases was also required to be >30. The primary INDEL readout was the number of INDEL events withing the repeat region per qualifying read. Given the stringency of full-length alignments, analyses were designed to detect small to moderate sized INDELs within the repeat region.

For the APOBEC3 mutagenesis analysis, reads were aligned to corresponding reference genomes. For A3B, A3B E255A, A3x3B, and A3x3B E255A samples, APOBEC analysis was performed on reads that exclusively mapped to the collapsed reference. For mEGFP reads were aligned directly to the mEGFP reference using the same full-length identity filters. The plus-strand reference sequence was scanned for 5′WGA motifs, corresponding to the canonical A3B-preferred substrate, 5′TCW, on the minus (cDNA) strand. A3B mutagenetic burden was measured as G-to-A substitutions in a 5′WGA motif (Phred quality >30). Mutation-load percentile plots were generated by pooling qualifying reads within each condition and ranking reads by WGA-context G-to-A mutations per read, splitting the ranked reads into 100 centiles, and plotting the mean mutation burden within each centile.

### QUANTIFICATION AND STATISTICAL ANALYSIS

#### Data analysis and figure generation

Illustrations of collapsing retroviral constructs were generated with BioRender. Data were analyzed and plotted in R (v4.5.0). Data was imported from Excel files using readxl, summarized using dplyr (https://CRAN.R-project.org/package=dplyr), and plotted using ggplot2 (https://ggplot2.tidyverse.org). Portions of the R scripts were drafted with assistance from

ChatGPT (OpenAI). All code was reviewed, modified, and validated by the authors before use. The Student's *t*-test was used to assess differences in viral recombination rate and APOBEC mutagenic burden. A threshold for significance was set at $P < 0.05$. Samples showing statistically significant differences relative to the indicated controls are annotated with *P*-values, and unlabeled comparisons did not reach the significance threshold.

### SUPPLEMENTAL INFORMATION

**Supplementary Figure 1.** Schematic of reverse transcription and second strand synthesis of CRV1.0

**Supplementary Figure 2.** Data from HeLa cells transduced with CRV2.0
